# Fusidic acid-based drug combinations exhibit enhanced activity against *Mycobacterium tuberculosis*

**DOI:** 10.1101/2023.01.19.524834

**Authors:** Charles Omollo, Atica Moosa, Kelly Chibale, Digby F. Warner

## Abstract

Tuberculosis (TB) imposes a major burden on global public health which is exacerbated by the escalating number of multidrug-resistant (MDR)-TB cases. There is consequently an urgent need for new anti-TB drugs and combination regimens. We have investigated the natural product antibiotic fusidic acid (FA) for repurposing against *Mycobacterium tuberculosis*, the causative agent of TB. Here, we report the results of synergy screens combining FA with a panel of approved anti-TB agents. Checkerboard and time-kill kinetics assays identified seven compounds from different chemical classes that synergized with FA in inhibiting the growth of *M. tuberculosis in vitro*: rifampicin (RIF), a rifamycin and frontline anti-TB drug; the macrolides, erythromycin (ERY), clarithromycin (CLR), and roxythromycin (ROX); the oxazolidinone, linezolid (LZD); the aminoglycoside, streptomycin (STR); and the aminocyclitol, spectinomycin (SPC). Among these, the strongest synergies were observed where FA was combined with SPC and ERY. Moreover, the FA-RIF combination was cidal, while all other FA combinations were bacteriostatic. These results provide *in vitro* evidence of the potential utility of FA-containing combinations against *M. tuberculosis*.

## INTRODUCTION

Combination therapy is essential to the clinical management of tuberculosis (TB) disease (1). Until recently, strategies to identify and advance promising combinations during early-stage pre-clinical TB drug discovery were limited. However, growing recognition of the need to identify new anti-TB drugs and regimens has re-focused attention on early-stage pre-clinical identification of synergizing combination partners for potential development (2), including drugs which are not clinically effective against TB (3, 4).

As part of a drug repurposing strategy, we utilized fusidic acid (FA) as anchor compound in developing matrix screening assays aimed at identifying optimal drug combination(s) that might be evaluated within the existing TB drug pipeline for potential clinical efficacy. FA, a translational inhibitor with demonstrated (albeit moderate) activity *in vitro* (5, 6), was selected owing to its unique mechanism of action: specifically, inhibition of mycobacterial protein synthesis by binding to elongation factor G (EF-G) (7). The antimicrobial-potentiating effect of FA with other antibiotics including the frontline anti-TB drug, ethambutol (EMB), as well as its lack of cross-resistance to other antimicrobial classes, provided additional motivation for our choice of FA (8, 9). In this short report, we present the analysis of *in vitro* interactions between FA and partner compounds comprising drugs currently used in TB treatment and selected translational inhibitors, the latter selected to enable evaluation the effects of combining FA with drugs acting on the same pathway **(Fig. 1)**, (10, 11).

**Figure 1.**
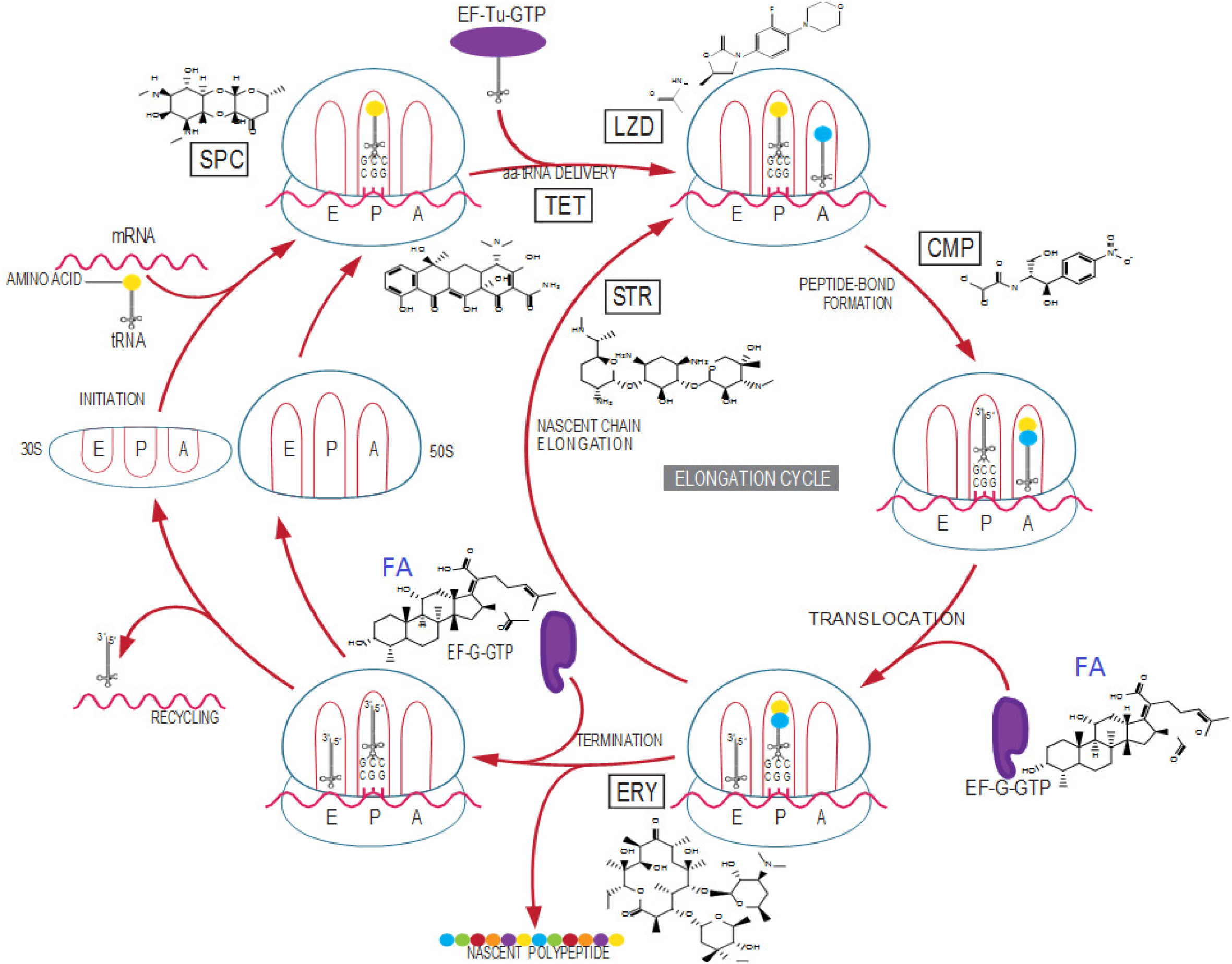
Antibiotic targets in protein synthesis: Schematic representation indicating known and predicted target sites of antibiotics disrupting different stages in the (myco)bacterial translation pathway. The tRNA binding sites - amino acid (A), peptide (P), and exit (E) - on the ribosome are indicated. Adopted and modified from Wilson *et al*. (10).

## RESULTS

### Checkerboard assay identifies synergistic drug combination partners for fusidic acid

To identify potential partners of FA, our preliminary screens utilized *Mycobacterium smegmatis* mc^2^155, a fast-growing, non-pathogenic mycobacterium which has been exploited as a useful surrogate in drug efficacy studies *in vitro*. In standard two-drug checkerboard experiments **(Table S1)**, SPC, ERY, CLR and TET exhibited synergy with FA, defined as FICI ≤ 0.5. These combinations displayed a 4-to 16-fold reduction in MIC_90_ for each of the individual drugs **(Fig. S1)**. No antagonistic effects were observed with any of the combinations tested.

The synergies detected in *M. smegmatis* informed subsequent selection of combination partners for evaluation in *M. tuberculosis* using the checkerboard method. In addition, representative drugs consisting of clinically used anti-TB agents (first- and second-line) and selected translational inhibitors were tested in combination with FA. For these assays, a *M. tuberculosis* H37Rv reporter strain expressing green fluorescent protein (GFP) was used, as described previously (12). **Fig. S2** provides an example of an interaction study in which FA and ERY were analysed in the checkerboard assay. Results in **Table 1** show similarity to those obtained for *M. smegmatis*, such that SPC, ERY and CLR exhibited synergy with FA against *M. tuberculosis*. The combination of FA and ERY returned a FICI value of 0.25, indicating a synergistic interaction **(Fig. S2A)**. Other drugs that synergized with FA included RIF, STR, roxithromycin (ROX), and LZD. These synergistic interactions generally resulted in 4-to 8-fold reductions in the MICs of each drug within the combination. Even though the combination of FA and BDQ did not result in a FICI value of ≤ 0.5, it is important to note that the two displayed approximately a 4-fold reduction in their respective MICs, and the observed FICI (0.55) was very close to that defined as “synergy”. No antagonistic interaction was observed between FA and any of the compounds tested. For the combinations exhibiting synergy with FA, isobolograms were constructed by plotting the FIC curves of the FA-drug combinations **(Fig. S3)**. Interactions between FA and ERY, SPC, and RIF were well within the synergy region (FICI < 0.5) whereas FA with STR, LZD, and ROX indicated borderline synergy (FICI = 0.5). The FA-INH interaction was included as a “no interaction” control.

**Table 1.**
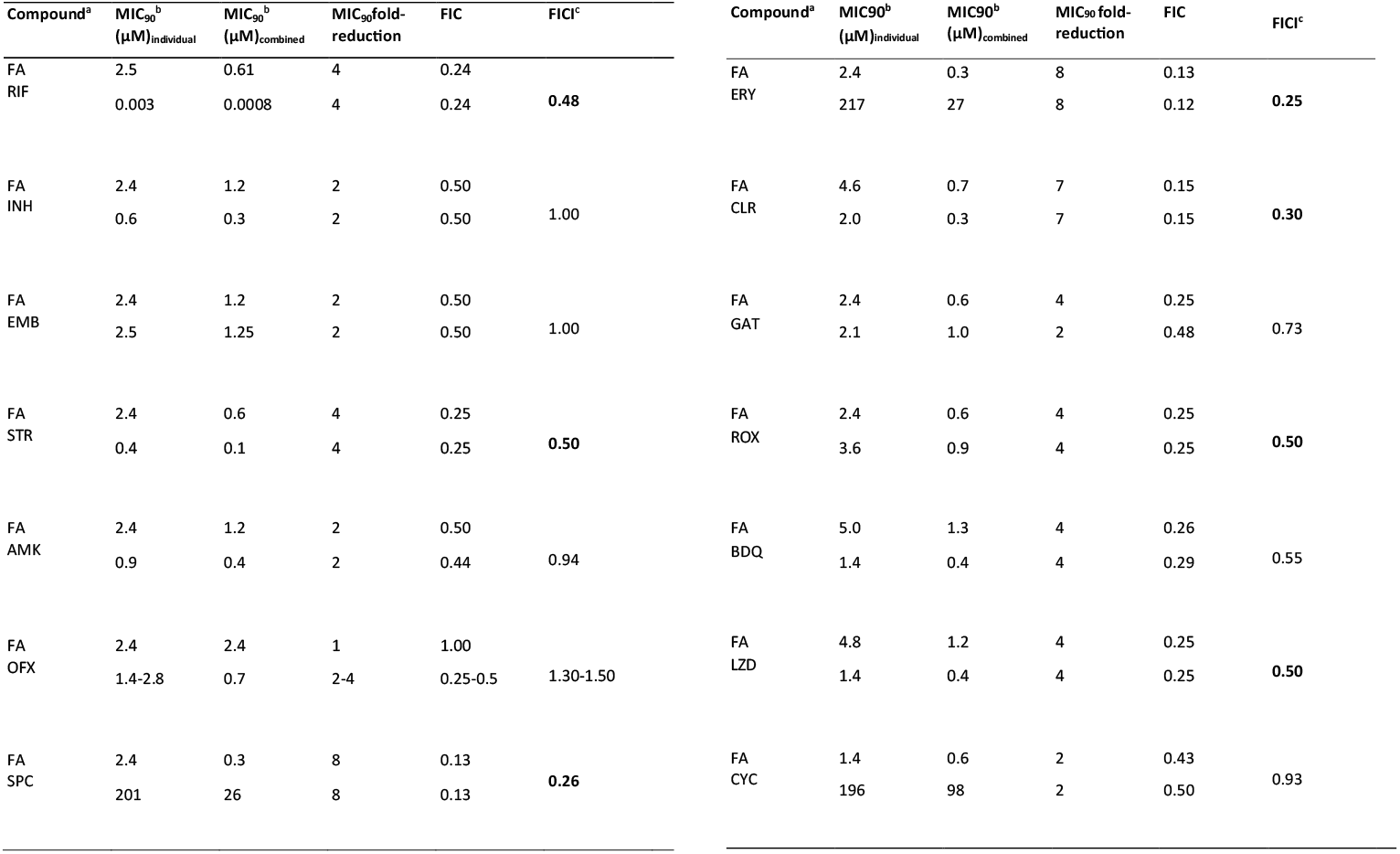
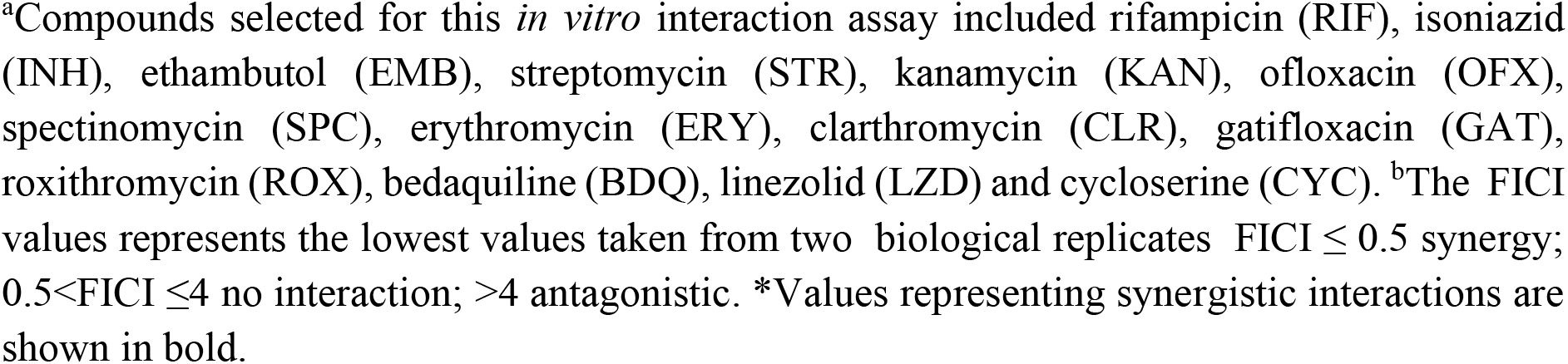
*In vitro* synergistic interaction between FA and anti TB agents or selected translational inhibitors against *M. tuberculosis*::gfp

To confirm results obtained using the checkerboard assay, the strongly synergistic FA-ERY combination was evaluated in a growth inhibition assay **(Fig. S4)**. For this purpose, FA and ERY were used at 0.3 and 27 µM, respectively, since these were the concentrations at which the lowest FICI value was obtained in the checkerboard assay **(Fig. S2)**. RIF, at a concentration of 0.015 µM, was included as a control. In the absence of drug, the population of *M. tuberculosis* increased over 14 days post-inoculation. In contrast, the population of viable cells remained relatively constant over the same duration when the growth medium contained the FA and ERY combination. Similarly, the medium containing FA, ERY plus sub-MIC RIF did not display any increase in the number of viable bacterial population over a 14-day period. In contrast, cultures incubated in the presence of the individual antibiotics, FA or ERY, showed equivalent growth to the untreated control.

### Assessing synergistic and peak plasma concentrations (C_max_) of FA synergizing drugs for optimal therapy

As a key consideration for the clinical potential of FA combinations, the respective concentrations at which synergy was observed were compared with the reported peak plasma concentrations (C_max_) for each drug. This is important in understanding whether the concentrations required to achieve the desired response are therapeutically feasible – and, consequently, whether the results obtained from an *in vitro* assay have any utility in guiding therapeutic use. Except for the FA-ERY interaction, synergies between FA and its partners were achieved at concentrations predicted to be below the maximum plasma concentrations (13, 14), suggesting that these concentrations are therapeutically achievable **(Fig. S5)**. For example, the presence of FA decreased the MIC of SPC from 201 µM to 3.14 µM, which represents a greater than 98% reduction in the MIC **(Table S2)**. This reduced concentration is far below the C_max_ value of SPC in humans (30.8 µM), determined following a 1000 mg intramuscular dose (15).

### Assessing cidal *versus* static synergies

To determine whether FA interactions resulted in killing or merely inhibited the growth of *M. tuberculosis*, the method of Zhang *et al*. was utilized (16). INH, a bactericidal agent, was used as a reference drug, and all drugs were tested alone and in combination against the *M. tuberculosis* H37Rv::*gfp* bioreporter strain. Results for the bacteriostatic or cidal effects of the drug combinations with FA are illustrated **(Figure 2)**. The FA-RIF combination displayed a MBC/MIC ratio of ≤ 2 on day 14 of evaluation, suggesting cidality. The other combinations tested – combining FA with SPC, ERY, CLR, ROX or LZD – all exhibited MBC/MIC ratios >2, implying static effects. The bacteriostatic/cidal action of individual drugs is shown in **Fig. S6**.

**Figure 2.**
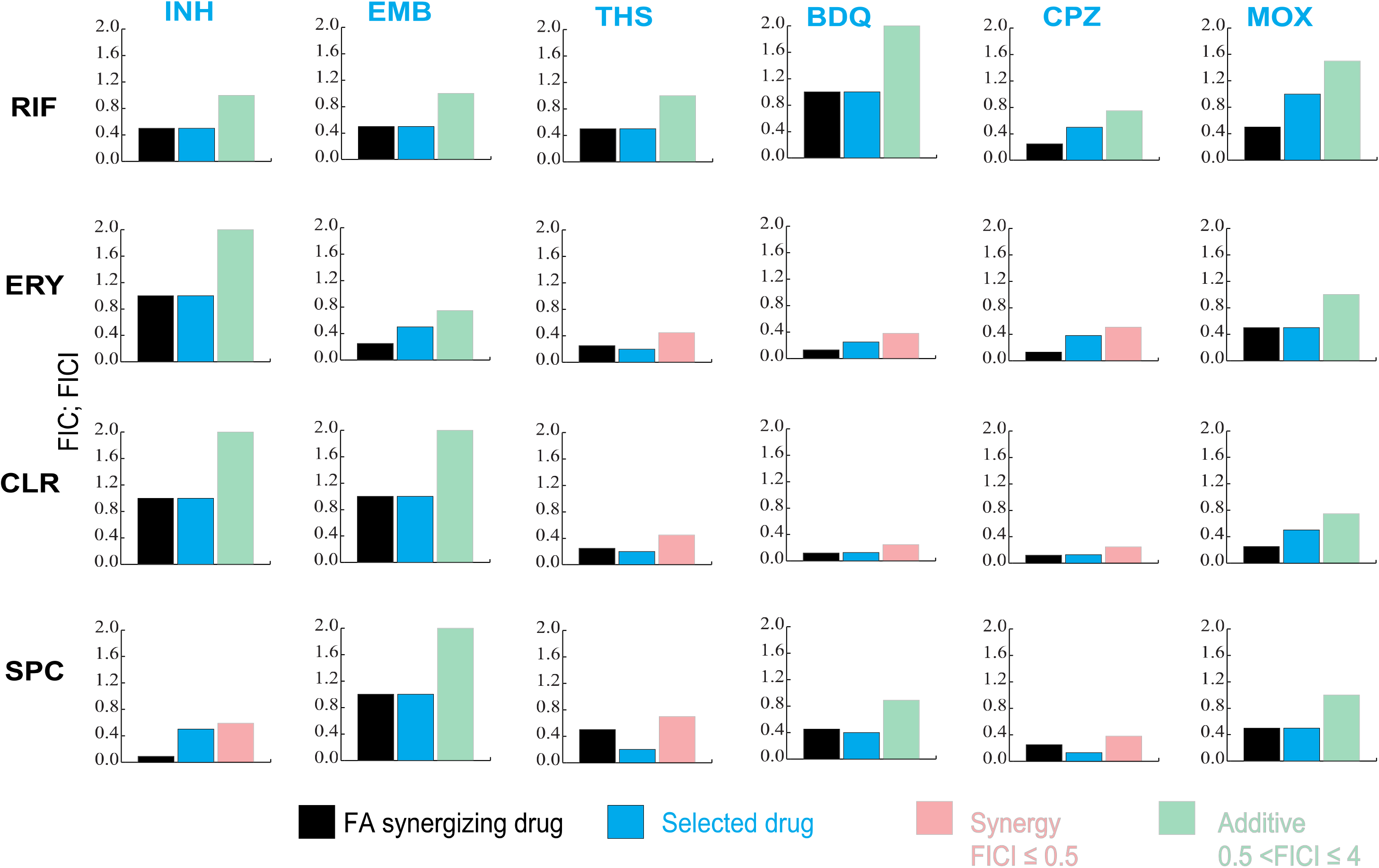
Bactericidal *versus* static effects of drug combinations against *M. tuberculosis*. Analysis of FA in combination with synergizing drugs; ERY, SPC, CLR, ROX, RIF, and LZD. INH was used as reference drug. MBC/MIC ratio ≤2: cidal; MBC/MIC ratio > 2: static. **\***Sub-inhibitory concentrations of FA that exhibited the respective synergies in these checkerboard experiments are between (0.3 – 1.2 µM) Data are from a representative experiment performed in triplicate. Error bars indicate standard deviations, calculated from the mean of triplicate samples.

### FA synergizing partners are active against the FA-resistant mutant

A cross-resistance study was performed using a FA-resistant *M. tuberculosis* mutant carrying H462Y mutation in *fusA1*(11). The FA-resistant mutant exhibited >100-fold MIC compared to the FA-susceptible parental strain. Six synergistic FA partner drugs – RIF, SPC, CLR, ERY, STR and ROX – were evaluated using the resazurin reduction microplate assay (REMA). The results **(Fig. S7)** indicated that the MIC_90_ values of these drugs remained the same against the FA-resistant mutant relative to the wild-type strain, confirming that there was no cross-resistance to each of the tested compounds.

### Interaction of FA synergizing partners with selected chemically unrelated compounds

Using a set of six chemically and structurally diverse drug agents **(Table S3)**, a combination assay was performed with four drugs that had displayed synergistic interaction with FA: RIF, ERY, CLR and SPC. The aim was to determine whether these compounds displayed synergy only with FA or exhibited a wider spectrum of synergistic interactions. The results of these pairwise combinations **(Figure 3)** revealed potentiating interactions between all FA synergizing partners (except RIF) and THS, BDQ and CPZ. CLR displayed strong synergies with BDQ and CPZ (FICI = 0.23 in both cases). The interaction between SPC and INH was also synergistic, while the remaining compounds (EMB and MOX) exhibited no interactions in combination with RIF, ERY, CLR and SPC.

**Figure 3.**
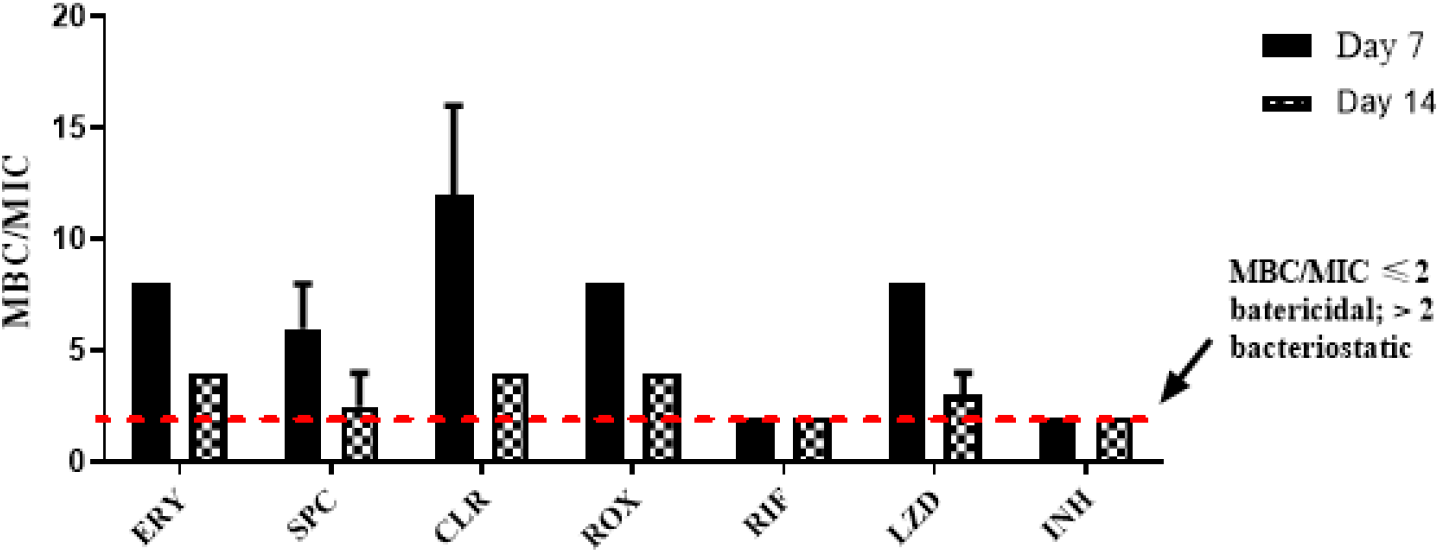
Drug interaction study. Pairwise interactions between selected drugs (blue fonts) and synergizing agents (black fonts) of FA (using *M. tuberculosis* H37Rv:*:gfp*. Red panels indicate synergistic interactions (FICI ≤ 0.5), while green panels are no interaction (0.5 < FICI ≤ 4). The bars indicate the fold-reduction in MIC required to achieve the same effect in combination *versus* single-drug assays. For example, in the interaction between RIF and MOX, top right-hand panel, the FIC for RIF = 0.45 (black bar) and MOX = 0.9 (blue bar). The resulting FICI for RIF-MOX = 1.35 (green bar) which indicates an additive interaction. Data are from the representative experiment performed in duplicate.

## DISCUSSION

Using a combination of checkerboard and time-kill kinetic assays, seven drugs were identified that synergized with FA *in vitro*. These include the rifamycin, RIF, and the macrolides, ERY, CLR, and ROX. Others included the oxazolidinone, LZD, the aminoglycoside, STR, and an aminocyclitol class compound, SPC. In all cases, the fold-reduction in the MIC_90_ of the interacting drugs was demonstrated; for example, ERY enhanced the potency of FA 8-fold, shifting the MIC_90_ from 2.4 µM to 0.3µM. All other two-drug combinations comprising FA plus a partner compound showed no interaction, with none exhibiting an antagonistic effect. Whereas some of these combination outcomes involving FA have been reported in previous studies (17), this study identifies new synergistic interactions by combining FA with LZD, ROX and STR, that can be exploited to guide rational efforts for development of effective combinatorial treatments.

Time-kill kinetics confirmed the strong synergy observed between FA and ERY. The mechanism underlying this interaction is not clear. However, precedent exists for the potency of combinations of compounds which disrupt the same cellular pathway or function (18). Both FA and ERY inhibit protein synthesis: ERY, like other macrolides, blocks the exit of the nascent peptide chain by binding to the 16S (19), while FA targets the EF-G elongation factor (20). Therefore, it’s possible that the combined inhibition of the translational machinery at separate processing stages underlies the observed whole-cell effect.

A key aspect to consider for the clinical utilization of (synergistic) drug combinations is whether the synergistic concentrations required for realizing an effective response are therapeutically achievable (21). Sufficient concentrations of the drug combinations in the bloodstream and infected tissues are needed to achieve the desired therapeutic effect (22). If both drugs are present with the concentration of one of the drugs being lower than the concentration for synergy, this can result in a mismatched exposure where the bacteria are exposed to long periods of monotherapy, risking development of genetic resistance (23). The results demonstrated that synergies with FA were achieved at concentrations lower than the respective peak plasma concentrations for all the drugs, except ERY. When FA synergizing partners were tested in combination with other chemically diverse agents, BDQ and CPZ exhibited the most promising synergistic interactions. The synergy with CPZ may be attributed to its efflux pump inhibitory effect, resulting in the accumulation of the partner compound (22). However, the mechanism responsible for synergistic interactions between BDQ and FA synergizing partners requires further investigation. A possible explanation for this synergy may be because of deleterious effects caused by combining FA with BDQ, the agent affecting energy metabolism.

The challenge of overcoming antibiotic resistance continues even with the use of drug combinations. Bacteria can evolve resistance to one drug thereby risking the possibility that they will also be able to resist the other drug(s) in the combination regimen. An understanding of cross-resistance between drugs is an important first-step in determining the drug combinations that would provide effective therapy (24). Here, the cross-resistance studies showed a mechanism of resistance specific to FA when compared to its synergizing partner compounds.

In summary, this work highlights promising synergistic interactions which might be investigated further against a broader panel of *M. tuberculosis* strains, including both drug- and multidrug-resistant clinical isolates.

### Materials and methods Chemicals and reagents

All chemicals and solvents were purchased from Sigma-Aldrich. Working solutions of all antimicrobial agents were prepared in dimethyl sulfoxide (DMSO).

### Bacterial strains and culture conditions

The laboratory strains, *Mycobacterium smegmatis* (*Msm*) mc^2^155, *Mycobacterium. tuberculosis* (*Mtb*) H37Rv and a reporter strain which has been used previously in high-throughput antimicrobial drug screening and constitutively expresses green fluorescent protein (GFP), H37Rv pMSP::eGFP (12), were maintained as freezer stocks. Strains were inoculated in standard Middlebrook 7H9 medium supplemented with 10% oleic acid-albumin-dextrose-catalase (OADC) (Difco) and incubated as stationary cultures at 37°C, sub-cultured, and incubated until culture density was approximately OD_600_ ∼0.5. Cell suspensions were diluted to give an expected final concentration of 10^5^ cells/ml at the time of inoculation into the microplate for the minimum inhibitory concentration (MIC) assays.

### Drug susceptibility assays

The Resazurin Microtiter Assay (REMA) was used to determine the susceptibility of drugs against *Msm* and *Mtb* strains, as described (25). Briefly, 2-fold serial dilutions of compounds were performed on clear, round-bottom 96-well plates using 7H9-OADC medium. Cultures grown to an OD_600_ of 0.5 (∼ 10^8^ cells/ml) and diluted 1,000-fold, a total volume of 100 µl per was added per well. The plates were sealed in zip-lock bags and incubated at 37 °C for 3 and 7 days: *Msm* and *Mtb* respectively, consistent with EUCAST guidelines (26) and published literature (27) recommending that MIC plates should be read after 7 and 14 days post inoculation. Resazurin dye was added, and the plates incubated for a further 24 h. Fluorescence readings, at excitation and emission wavelengths of 540 and 590 nm respectively, were recorded using a BMG Labtech POLARstar Omega plate reader (BMG Labtech, Offenburg, Germany) or a SpectraMax i3x plate reader (Molecular Devices). The lowest drug concentration which inhibited growth by at least 90% was determined from the dose-response curve; this concentration was defined as the MIC_90_ value.

### Checkerboard assays

#### 2D Checkerboard

Standard “two-dimensional” (2D) drug-drug interactions were determined by checkerboard titration in a 96-well plate (11). The 2D microdilution was carried out as described (28), with slight modification. Briefly, the first drug (A) to be serially diluted was dispensed (2 µl) along the x-axis (columns 3 to 11, row B) at a starting concentration 100 times higher than the final concentration in the well, and (2 µl) per well of the second drug (B) was serially dispensed along the y-axis (from row B to H) at a starting concentration 100 times higher than the final concentration in the 96-well microtitre plate. The first column (column 1) and last column (column 12) contained drug-free controls (with 1% DMSO as a diluent) and a control drug concentration giving maximum inhibition, respectively. The second column from B2-H2 and first row from A3-A11 contained individual drugs, thus providing the MIC for each drug alone in each assay (each plate). The plates were placed in zip-lock bags and incubated at 37 °C for 7 days. Resazurin dye was then added and the plates incubated for a further 24 h. A visible colour change from blue to pink indicated growth of bacteria, and the visualized MIC was defined as the lowest concentration of drug that prevented growth (at which the colour change did not occur) (25). Fluorescence readings (excitation: 544 nm, emission: 590 nm) were obtained using a BMG Labtech POLARstar Omega plate reader (BMG Labtech, Offenburg, Germany) or a SpectraMax i3x plate reader (Molecular Devices). The mean fluorescence value for the “maximum inhibition” wells (column 12) was subtracted from all other wells to control for background fluorescence. Percent inhibition was defined as 1-(test well fluorescence units/mean fluorescence units of maximum inhibition wells) × 100 on day 8 after incubation. The lowest drug concentration effecting inhibition of 90% was considered the MIC_90._ In addition, synergy was interpreted according to the sum of fractional inhibitory concentration (∑FIC). The fractional inhibitory concentration for each compound was calculated as follows (29): FIC_A_ _ (MIC of compound A in the presence of compound B)/(MIC of compound A alone), where FIC_A_ is the fractional inhibitory concentration of compound A. Similarly, the FIC for compound B was calculated. The ∑FIC was calculated as FIC_A_ + FIC_B_. Synergy was defined by values of ∑FIC ≤ 0.5, antagonism by ∑FIC > 4.0, and no interaction by ∑FIC values from 0.5 to 4.0 (30).

#### CFU enumeration

To estimate the number of live bacilli after drug treatment, untreated and drug treated *Mtb* cells were plated onto 7H10 agar. Colonies were counted after 3-4 weeks of incubation at 37°C.

### Bactericidal versus bacteriostatic analysis

In this assay, an established technique was used to differentiate the bacteriostatic or bactericidal effect of a single drug or drug combination (16). Briefly, the MIC_90_ of the drugs against *M. tuberculosis* H37Rv::gfp or wild-type strain were first determined in 7H9 broth using the standard microdilution method on a 96-well plate. Briefly, 2-µl of the test drugs were added to the 7H9 medium to make a total of 100µl. This was followed by 2-fold dilutions on a 96-well plate. 50-µl of mid-log phase culture (OD_600_ 0.6) adjusted to approximately 10^5^ CFU/ml were added to the wells. The plates are then packed in zip-lock bags and incubated at 37°C for 7 days. Thereafter, the MIC_90_ was established by measuring fluorescence using the BD microplate reader with excitation and emission wavelengths of 485nm and 520nm, respectively. 5-µl from every well was later transferred to 100-µl of a drug-free medium in another 96-well plate, MBC plate, and incubated at 37°C for a further 7 days. Fluorescence was measured on day 7. The concentration at which there was 90% or greater inhibition of RFU compared to those of untreated cells was defined as the bactericidal concentration. A compound or drug combination was considered bactericidal if the ratio between the minimal bactericidal concentration MBC_99_ and the MIC_90_ was equal to or smaller than 2, that is MBC/MIC ≤ 2. Alternatively, when the wild-type strain was utilized, the transfer of 5-µl was effected before adding 10-µl per well of resazurin blue, this was then incubated for 24 h. A change from blue to pink indicated growth of bacteria, and the MIC was defined as the lowest concentration of drug that prevented this colour change. INH, a bactericidal drug was used as a positive control.

### Statistical Analyses

Statistical analyses were performed using Prism 9.0.0.121 (GraphPad). Means were compared via ANOVA, with post-test evaluation using Dunnett’s or Bonferroni’s test.

## ACKNOWLEDGEMENTS

The authors are grateful for the funding support of the South African Medical Research Council. KC is the Neville Isdell Chair in African-centric Drug Discovery and Development and thanks Neville Isdell for generously funding the Chair. The South African Research Chairs Initiative of the Department of Science and Innovation is also gratefully acknowledged for their support (KC).

